# Non-inertial scan angle multiplier for expanded fields-of-view

**DOI:** 10.1101/2025.06.13.659647

**Authors:** Che-Hang Yu, Yiyi Yu, Joseph S. Canzano, Yuandong Fei, Spencer LaVere Smith

## Abstract

In many laser scan engines, mechanical inertia imposes a tradeoff between mirror diameter (mass) and scan speed (cycle time) and/or scan angle amplitude. Ultimately, this limits the field-of-view and/or scan speed in multiphoton and confocal imaging, laser-based manufacturing, and other applications. To push parameters past this inertia limit, we present a non-inertial (stationary) add-on unit that doubles the scan angle amplitude while preserving both the beam size and cycle time, thus doubling the étendue (optical invariant) and overall system throughput. We demonstrate its use in two-photon calcium imaging of neural activity in living mice. We also adapt the approach to present a phase doubling unit that doubles the maximum range of phase modulators. Finally, we describe further variants including generalizing the approach to Nx scan angle multiplication (N = 2, 3, 4, …) with diffraction limited performance across a wide scan angle range and over a broad wavelength bandwidth. These non-inertial scan angle multipliers (called Nisam2x, 3x, …) can expand the capabilities of a range of technologies including imaging, adaptive optics, sensing, marking, and manufacturing.

## 1. Introduction

Laser scan engines are used in a broad array of technologies, including imaging, sensing, marking, and manufacturing. In many of these applications, the speed of scanning and field-of-view are key figures-of-merit. These figures of merit can be limited by mechanical inertia when moving mirrors are used to scan the beam. Non-inertial scan systems such as acousto-optical deflectors can provide very rapid scanning, but typically over relatively small angles and with small beams compared to moving mirrors. For larger beams, such as those used in high numerical aperture imaging, moving mirrors offer the largest ranges of angles. This aspect of laser scanning can be characterized using the concept of the optical invariant, étendue, or optical throughput [1, 2].

Galvanometer mirrors are commonly used in laser scanning. Their scan cycle frequency and scan angle amplitude determine the scan speed and the field-of-view, respectively. Closed-loop scanners (also known as servo, or linear scanners) can follow arbitrary command waveforms, while resonant scanners ue a mechanical resonance to offer higher scan speeds, but in a free-running, open-loop mode. A major figure of merit for scan mirrors and scan engines is the maximum scan amplitude, which can be specified as either a peak-to-peak value or a plus-minus (±) range, which is simply half of the full peak-to-peak scan amplitude. Scan ranges can be either mechanical or optical. The mechanical scan range is the actual mirror rotation, and the optical scan range is the angle deflection exhibited by a scanned laser beam (**Fig. 1a**). The optical scan range is simply two times the mechanical scan range. We mostly use the optical scan range in this report. The scan frequency is ultimately limited by mechanical inertia (**Fig. 1b**). Thus smaller mirrors (i.e., beam diameters) and/or smaller scan amplitudes are used to reach higher scan frequencies. Due to the trade-offs between the scan frequency and the scan angle of resonant scan mirrors, there is a figure of merit called the optical invariant (or étendue), for example, as the product of the beam radius and the scan amplitude [1].

**Fig. 1.**
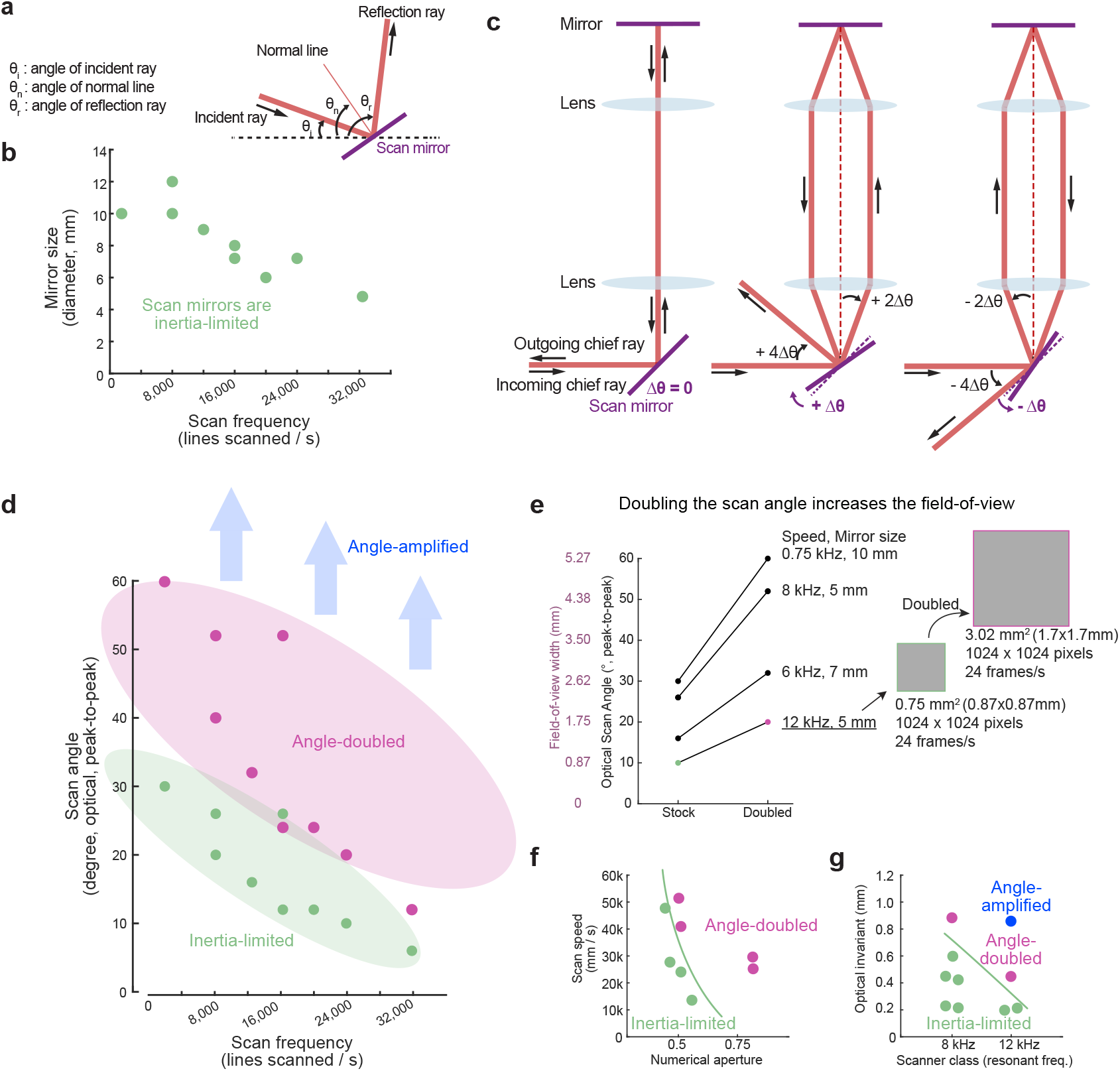
Overcoming inertia limits with scan angle doubling and amplification. (a) A schematic shows the definitions of notations for scan mirror reflection. (b) Resonant scanning mirrors show decreasing scan angles as their scan frequencies increases due to inertia limits. (c) The optical layout of an angle doubler, illustrating the operating principle. (d) Conventional scanners are limited by inertia to trade off scan angle with scan frequency (not to mention mirror size as noted in panel (b)). Now, with angle doubling and angle amplifying, the scan angle amplitude can be increased with no trade-offs in scan frequency (or mirror size), unlocking more of the parameter space for system design. (e) The scan angle and the corresponding field-of-view of common commercial resonant scanners before and after angle doubling when a 10x objective lens is used in a laser scanning microscope. The imaging area is increased by 4 fold with angle doubling while the imaging speed (line rate) and the mirror size remain the same. (f) Angle doubling unlocks parameter space for design in terms of scan speed (mm of sample plane scanned per second) versus numerical aperture (resolution), enabling high speed, high resolution laser scanning. (g) Angle-doubling and angle amplification enable higher optical invariant scan engines with no trade-off in scan speed. The 4x amplification of scan amplitude for a 12 kHz scanner is plotted in the top right.

We consider an example: laser scanning multiphoton microscopy [3, 4]. Many applications benefit from micron resolution over millimeter length scales and subsecond time resolution [1,5–9].

The microscope objectives for high resolution, large field-of-view systems are physically large. This is necessary to achieve a large field-of-view (long effective focal length) while maintaining a high optical resolution (numerical aperture). This physical size is related to a large optical étendue or optical invariant. This value can be defined as the product of the field-of-view radius and numerical aperture, or as the product of the aperture radius and the sine of the maximal acceptance angle of incidence at the objectives’ back aperture [1, 5].

To fully use the optical invariant of the objective, the scan engine must deliver a beam with a diameter and scan angle range matched to the objective [1, 2, 10, 11]. For slow scanning, this is feasible without exotic instrumentation. However, high speed scanning with large diameter beams suffers from the tradeoffs described above. Some systems provide resonant scanning across the full field-of-view, but typically at a lower NA and field-of-view size [5, 12]. The systems with the largest high resolution fields-of-view provide resonant scanning across only a portion of their field-of-view. These “resonant-galvo-galvo” systems include a third mirror. In addition to the slow Y-axis mirror, and the resonant X-axis mirror, another slow servo galvanometer is relayed into the scan engine to provide additional deflection angle for the X-axis scanned by the resonant mirror. Thus, these systems image resonant-scanned “stripes” across the field-of-view. For example, the 2p-RAM mesoscope is equipped with a 12 kHz resonant scanner, which spans only 0.6 mm of its 5 mm wide field-of-view [7]. Similarly, the DIESEL2p system is equipped with an 8 kHz resonant scanner that spans 1.5 mm along the resonant axis, which again is smaller than its 5 mm wide field-of-view [6].

One can imagine synchronizing two mirrors and optically relaying the beam to double the scan range. Instead of using two mirrors, a stationary mirror could be used to reflect the beam back to the scanner. Thus the beam impinges twice onto the same mirror before propagating to the other optics (e.g., scan lens). This idea has been explored with a polygonal scanner [13], where a 1.7x increase in the scan angle was obtained, and in a microlens array study [14] which used the arrangement to generate multiple scan spots. Here we present a general approach for scan angle doubling, where the optical performance achieves full doubling of the scan amplitude, and does so without added optical aberrations, and without trading off mirror / beam diameter or scan frequency. The results we present here demonstrate what is, to our knowledge, the largest optical invariant scan engine designs for resonant scan mirrors to date. In addition, we present some alternative embodiments including a phase doubling approach and a way to increase the scan angle by multiples greater than two. Thus, we show that the optical invariant can be doubled or tripled or more, and therefore by induction, the scan angle can be independently engineered separate from the scan frequency, without limitations from inertia.

## 2. Results

### 2.1 Principle of operation

**Figure 1c** shows a schematic representation of an angle-doubling unit. It is a four focal length (4-f) optical relay, consisting of a scan mirror, a pair of lenses, and a flat mirror perpendicular to the optical axis of the 4-f relay. When the scan mirror is at its neutral scan angle (Δ*θ* = 0), an incident ray reflected by the scan mirror travels through the center line of the 4-f relay, gets reflected off the flat mirror, and returns through the same 4-f relay along the same path of the incoming ray **(Fig. 1c, left)**. As the mirror rotates clockwise by +Δ*θ* **(Fig. 1c, middle)**, the incoming ray is reflected off the scan mirror by +2Δ*θ*, and gets refracted by a pair of lenses to reach the flat mirror. After the ray hits the flat mirror, it returns along a route that is mirror-symmetric to the center line (black dotted line) of the 4-f relay this time, and hits the scan mirror again. Because the ray is not returning along the same route, the outgoing ray is not overlapping with the incoming ray. Instead, the ray exits at an angle of +4Δ*θ* relative to the incoming ray. Following the same rule, as the scan mirror rotates counter-clockwise by –Δ*θ*, the final outgoing ray is reflected by −4Δ*θ* relative to the incoming ray **(Fig. 1c, right)**. As a result, the outgoing ray scans over a peak-to-peak optical range of ±4Δ*θ* while the scan mirror scans ±Δ*θ* mechanically. This result is in contrast to the normal use case where a scan mirror with a scan range of ±Δ*θ* only offers a ±2Δ*θ* peak-to-peak scan angle optically. Therefore, the method we presented doubles the scan angle from ±2Δ*θ* to ±4Δ*θ*, and the doubling process is independent of, thus decoupled from, the scan frequency and the scanner size (aperture or mass). In other words, we have doubled the étendue or optical invariant, as measured as the area of the entrance pupil times the solid angle the source subtends as seen from the pupil, unlocking a broader array of the parameter space for design **(Fig. 1d-g)**.

The angle doubling can be described mathematically, where *θ*_*i*_, *θ*_*r*_, *θ*_*n*_ are angles of incidence, reflection, and the normal vector from the surface of the mirror, all measured from a reference horizontal line. For mirror reflection, the reflection angle is equal to the incident angle **Fig. 1a**:

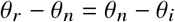

Knowing the *θ*_*i*_ and *θ*_*n*_, *θ*_*r*_ can be calculated as:

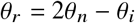

In the angle doubling system shown in **Fig. 1c**, the light interacts with the scanning mirror two times. For the first reflection, the incident angle of the light is *θ*_*i*_=0, and the normal line of the mirror is scanning over a range of *θ*_*n*_ = 45°± Δ*θ*, and the angle of reflection, *θ*_*r*_, can be computed as:

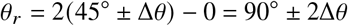

When the light returns to the scanning mirror, using the prime mark ‘to indicate the angles on the return trip, the new incident angle is 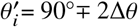, while the angle of the mirror normal line stays the same as 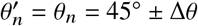. Hence, the second angle of reflection, 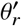, is:

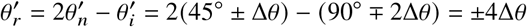

This derivation shows that the scan angle of the light is ±4Δ*θ* when the scanning mirror scans over a range of ±Δ*θ* mechanically in an angle doubling system. The range ±4Δ*θ* is two times larger than the scan range without the angle doubling system. The slow axis can be increased arbitrarily because the slow mirror is not typically inertia-limited. Thus, a four-fold larger field-of-view laser scanning system can be achieved while maintaining the original beam diameter and line scan rate (**Fig. 1e**).

### 2.2 Experimental validation

To validate the concept, we built a simple angle-doubling unit **(Fig. 2)**. A laser was focused onto a conjugate plane, passed through a collimating lens onto a scanning mirror, reflected to another lens, and come to focus at a second conjugate plane. Then the beam continued through another collimating lens, reflected back off of a stationary mirror, and retraced a new path through the elements to exit the system. At the conjugate planes, the scan angles were converted to scan lines as the focus translated across the scan axis. The length of the scanned lines was linearly proportional to the scan angle magnitude. To measure these scan lines, we picked off a bit of power from the conjugate planes using glass slides (dashed orange lines in **Fig. 2a**), and projected them onto paper rulers (yellow lines in **Fig. 2a** and inset **Fig. 2b**). As the scan mirror scanned at ±5 degrees optically, we found that the length of the line from after the angle-doubling unit (∼33 mm) was two times longer than that of the line from before the angle doubling unit (∼17 mm) **(Fig. 2b)**. This experimental result of a two-fold difference in the scan angle validated our proposed method for angle doubling.

**Fig. 2.**
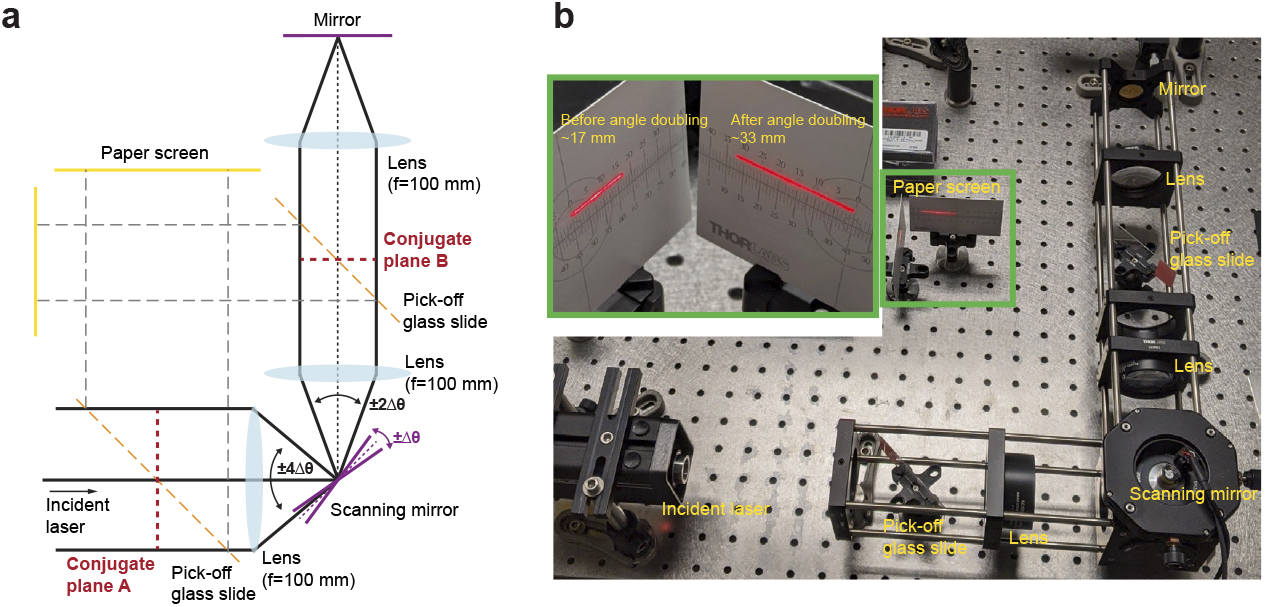
Experimental measurement of an angle doubling system. (a) A schematic diagram shows the experimental setup of an angle doubling setup built with off-the-shelf lenses (focal length = 100 mm, LA-1050-A, Thorlabs), mirrors, and a galvo scan mirror. (b) A photograph shows the actual setup of an angle doubling system. The inset is a view of the paper screens that show the amplitude of the scanning laser projected from the conjugate planes A and B while the galvo mirror scans at ± 5° optically.

### 2.3 Angle doubling for two-photon imaging

Next, we sought to confirm that the approach could provide high performance in a demanding application. The fastest resonant scanners, e.g., 12 kHz, have relatively small maximal scan amplitudes. While their high speed is desirable, their low scan angles result in reduced fields-of-view. So we integrated an angle-doubling unit into a laser scanning two-photon microscope to overcome this limitation. The system consists of two perpendicular arms, with a 4-f relay in each arm **(Fig. 3a)**. The angle-doubling arm consists of two identical scan lenses (LSM54-1050, Thorlabs), with effective focal lengths (EFL) of 54 mm. The fast axis was scanned with a 12-kHz resonant scan mirror (CRS 12 KHz, Cambridge Technology), which is limited to a ±5° peak-to-peak optical scan angle. Instead of a stationary mirror in the relay for angle doubling, we used a linear galvanometer scan mirror (±20° scan angle maximally; 6220H, Cambridge Technology) to scan the slow axis. The other 4-f arm was the two-photon imaging arm, consisting of a scan lens (EFL = 54 mm, LSM54-1050, Thorlabs) and a tube lens (EFL = 200 mm, TTL200MP, Thorlabs) followed by an objective lens. The beam magnification is 200/54 = 3.7x and the scan angle is minified by the reciprocal factor, 0.27x. These two arms were configured so that the resonant scan mirror and the linear galvanometer scan mirror were conjugated to the back aperture of the objective lens.

**Fig. 3.**
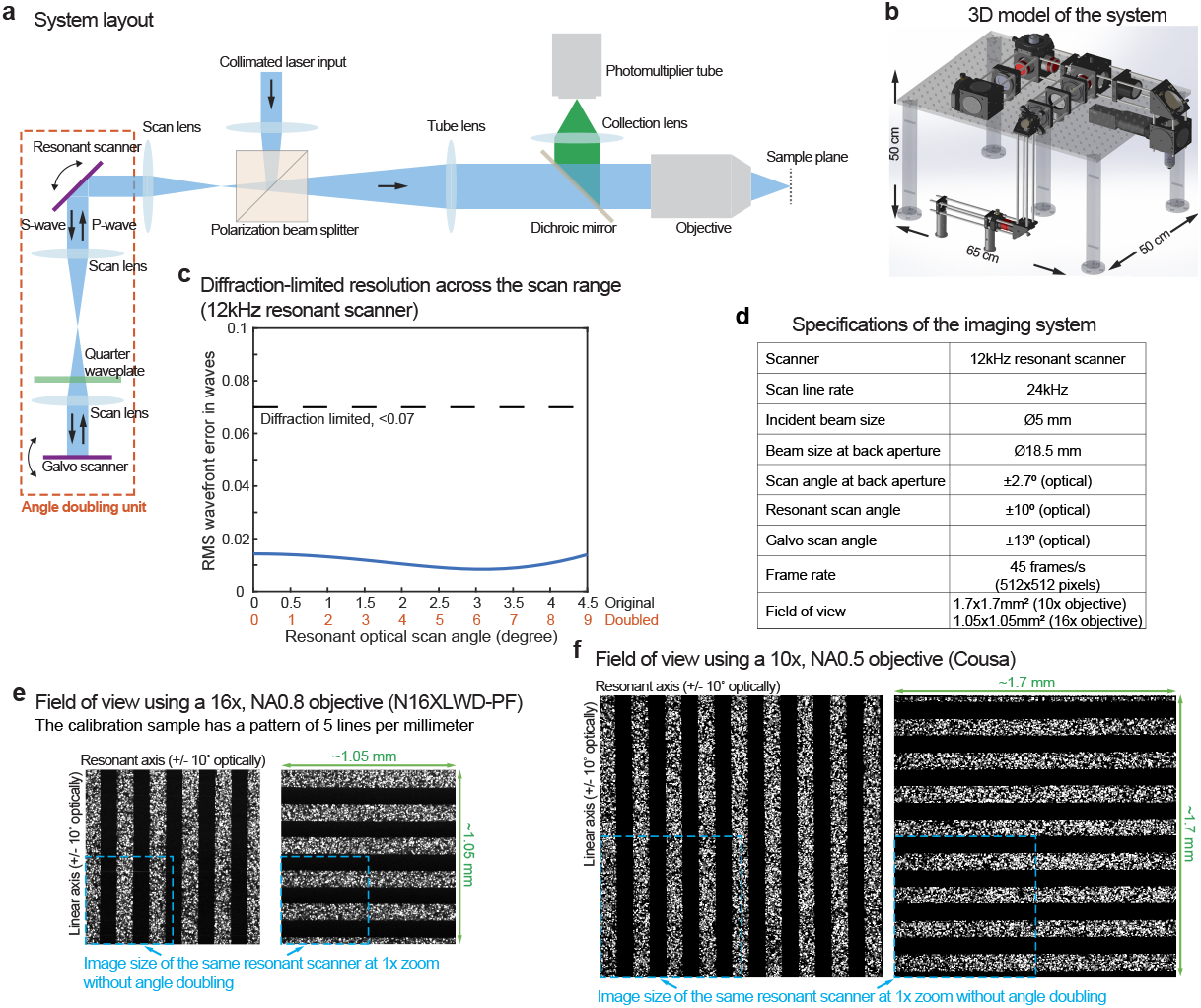
Integration of an angle doubling unit with a 12 kHz resonant scanner into a two-photon microscope. (a) A schematic diagram shows the integration of an angle doubling unit with a two-photon microscope. (b) A 3D model of the full optomechanical setup, including the footprint dimensions. The resonant mirror has a scan frequency of 12 kHz and an optical scan angle of ±5°. After angle doubling, the resultant scan angle is ±10° optically. (c) An optical simulation of (b) shows the root-mean-square error of the wavefront as a function of the scan angle of the resonant scanner at the imaging plane. (d) A specification table lists the parameters of the imaging system with the angle-doubled 12 kHz resonant scanner. (e, f) Images of a fluorescent ruler (5 lines/mm) were acquired using a Nikon 16x/NA0.8 water-immersion objective (e) and a Cousa 10x / 0.5 NA air objective (f). The fluorescence ruler was imaged in vertical and horizontal orientations to calibrate the field of view in the resonant x-axis and galvo y-axis. With angle doubling, the field of view is 1.05 × 1.05 mm^2^ with the 16x objective (e) and 1.7 × 1.7 mm^2^ with the 10x objective (f) at 1x zoom; blue dashed squares indicate the field of view without angle doubling at 1x zoom.

Polarization optics are used to direct the laser through the system. The input beam was fully S-polarized and reflected by the polarizing beam splitter (PBS; PBS513, Thorlabs) to direct it to the angle-doubling unit. The polarization beam splitter cube introduces dispersion, which could be mitigated by using a polarizing beam plate. To couple the angle-doubled beam into the two-photon imaging arm, a quarter-wave waveplate (39-046, Edmund Optics) was inserted in the angle-doubling arm. After passing through the quarter-wave waveplate twice, the polarization of the laser beam was fully P-polarized and passed through the PBS to reach the imaging plane under the objective.

The signal generated from the imaging plane was collected by the objective and directed to the photomultiplier tube (PMT2102, Thorlabs) via the dichroic mirror (DI03-R785-T1-50.8×50.8, AVR Optics) and the collection lens (LA1050-A-ML and ACL2520U-A,Thorlabs). The dimension of the system were, in cm: 65 × 50 × 50 (length x width x height) **(Fig. 3b)**. We modeled the full system with raytracing software (OpticStudio, Ansys Zemax) to evaluate the optical performance. The root-mean-square error of the wavefront was well below the criterion of 0.072 wavelengths, confirming diffraction-limited resolution across the accessible scan range **(Fig. 3c)**. The specification table summarizes the detailed parameters of this imaging system **(Fig. 3d)**.

We verified the angle doubling by measuring the field-of-view acquired, using a fluorescence sample with a periodic structure (5 lines per millimeter). With a 16x 0.8 NA water immersion objective (EFL = 12.5 mm, Nikon), and the fast axis scanning ±5°, the field-of-view was 1.05 × 1.05 mm^2^. As noted above, in this scan engine, the scan angle was minified by 0.27x, so at the objective the scan angle was 1.35 degrees. The field-of-view without doubling should have been 2 * 12.5 mm * tan 1.35° = 0.58 mm. Once doubled, and multiplied by the 90% fill fraction used in the scan control software (ScanImage, MBF) we get 0.58 * 2 * 0.9 = 1.04. Thus, the 1.05 mm wide field-of-view represents a doubling of the field-of-view. Similarly, when a 10x 0.5 NA air objective (EFL = 20 mm, Cousa [15]) was used, the field-of-view was 1.7 × 1.7 mm^2^ **(Fig. 3f)**, again matching the prediction (before doubling: 2 * 20 mm * tan(1.35°) = 0.94 mm; after doubling and 90% fill fraction: 0.94 * 2 * 0.9 = 1.7 mm).

The angle doubling does not require any imaging trade-offs. The scan frequency of the resonant scan mirror is the same after doubling, the scanning is simply spanning twice the distance in the image plane as it would without doubling. Moreover, the beam size at the back pupil of the objective was the same, preserving the NA and resolution of imaging. Together, these results demonstrated the compatibility of the angle doubling system with two-photon microscopy, and how it can increase the field-of-view with no trade-offs.

#### 2.3.1 In vivo imaging with angle-doubling at 24,000 lines/s scanning

To demonstrate the data quality with angle-doubling, we performed *in vivo* two-photon [3, 4] calcium imaging in genetically engineered mice, expressing the genetically encoded calcium indicator, GCaMP6s [16, 17]. This fluorescent calcium indicator is used to image neural activity in large populations of neurons [16]. The cranial window was surgically implanted over visual cortex (V1) as previously detailed [18]. The imaging field of view with a 16x objective (0.8 NA; Nikon) is 1.05 × 1.05 mm^2^ (512 × 512 pixels) and a frame rate of 45 frames / s. Neurons are clearly resolved **(Fig. 4a; Visualization 1)**, and calcium activity from at least 243 neurons were detected **(Fig. 4b and c)**, showing the fidelity of the signals in the obtained imaging data. We can further increase the size of the field-of-view by using a lower magnification objective. Using a 10x (0.5 NA, 20 mm working distance [15]), the imaging field-of-view is 1.7 × 1.7 mm^2^ (1536 × 1536 pixels) at an imaging rate of 15.6 frames / s, and at least 1507 active neurons were detected **(Fig. 4e-f; Visualization 2)**. In contrast, the image area in a square raster scan at 1× zoom without angle doubling is reduced to one-quarter (25%) of that achieved with angle doubling. Therefore, our angle-doubling approach can quadruple the field-of-view of two-photon imaging while maintaining the same high resolution and scan frequency offered by the 12-kHz resonant scan mirror.

**Fig. 4.**
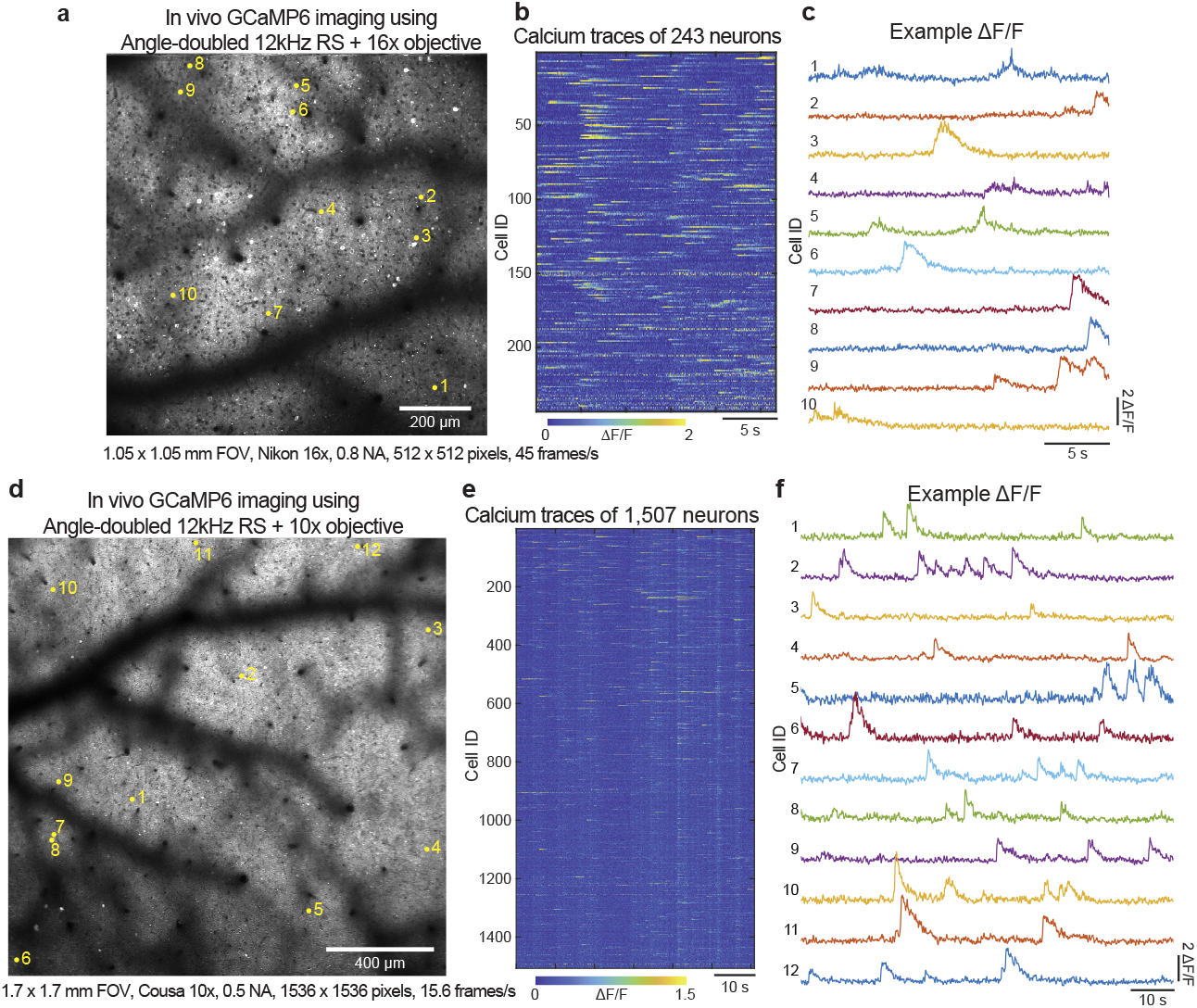
*In vivo* imaging of calcium activity in a mouse brain using angle-doubled 12kHz resonant scanner with different objectives. (a) Population calcium imaging (GCaMP6s) over a 1.05-mm-wide FOV at 45 frames/s using the angle-doubled 12kHz resonant scanner with the Nikon 16x objective (NA 0.8). See also the associated **Visualization 1**. (b) 243 neurons were identified and their calcium traces were shown. (c) Calcium traces from neurons labeled in (a) were plotted. (d) Population calcium imaging (GCaMP6s) over a 1.7-mm-wide FOV at 15.6 frames/s using the angle-doubled 12kHz resonant scanner with the Cousa 10x objective (NA 0.5). See also the associated **Visualization 2**. (e) 1507 neurons were identified and their calcium traces were shown. (f) Calcium traces from neurons labeled in (d) were plotted.

#### 2.3.2 In vivo imaging with angle-doubling and large étendue scan optics

The angle doubling technique can be readily applied to other resonant scanners, and the scan range increase can surpass the étendue of conventional scan optics, requiring new optical designs. We implemented angle doubling on an 8 kHz resonant scanner to further expand the field-of-view for two-photon imaging taking the advantage of its original ±13° scan amplitude. Doubling the scan angle from ±13° to ±26° without altering the mirror size and scan frequency doubles the optical étendue, exceeding the limits of off-the-shelf scan and tube lenses, which lack the aperture size to accommodate such large angles, resulting in severe vignetting **(Fig. 5a)**. Consequently, new large étendue scan optics were required. The system is configured to deliver a 20-mm beam diameter and ±5° scan at the objective’s back aperture. To achieve this, we used a new 50-mm effective focal length scan lens that supports a 5-mm beam at ±20° scan (Ventana SL; Pacific Optica), followed by a 200-mm effective focal length tube lens (Ventana TL; Pacific Optica), which expands the beam to 20 mm and ±5° at the objective’s back aperture. The resulting scan engine features broad achromaticity and anti-reflection coatings across 800–1350 nm, and supports low root-mean-square wavefront error and a high Strehl ratio throughout the field-of-view, facilitating simultaneous two-photon imaging, three-photon imaging, and two-photon optostimulation **(Fig. 5a-e)**.

**Fig. 5.**
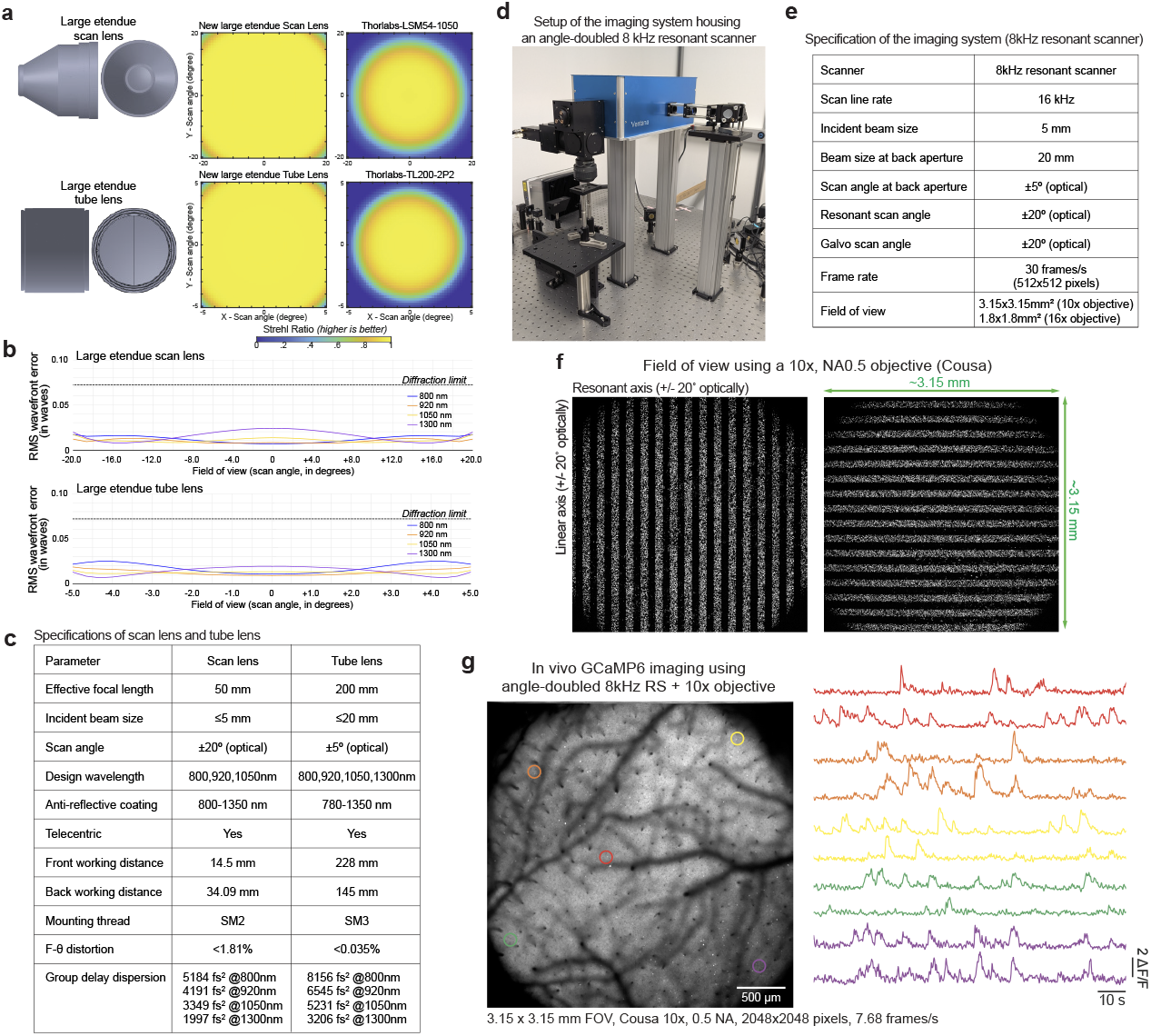
*In vivo* calcium imaging in a mouse brain using angle-doubled 8kHz resonant scanner and custom large étendue optics. (a) 3D rendering of the newly developed large étendue scan lens and tube lens. The middle column shows 2D Strehl ratio maps as a function of scan angle, while the third column presents comparable data for off-the-shelf optics. (b) Root-mean-square (RMS) wavefront error plots for the scan lens and tube lens at 800, 920, 1050, and 1300 nm, demonstrating low wavefront errors and a wide margin from the diffraction limit across large scan angles. (c) Specification table detailing parameters of the scan and tube lenses. (d) Photograph of the custom two-photon imaging system integrating the angle-doubled 8kHz resonant scanner with the large étendue optics. (e) Specification table listing parameters of the imaging system in (d). (f) Fluorescent ruler images (5 lines/mm) acquired with a Cousa 10x/NA0.5 air objective. The ruler was imaged in vertical and horizontal orientations to calibrate field of view in the resonant x-axis (±20°) and galvo y-axis (±20°). With angle doubling, the field of view is 3.15 × 3.15 mm^2^ at 1× zoom with no stitching needed. (g) Population calcium imaging (GCaMP6s) over a 3.15-mm-wide field of view at 7.68 frames/s using the angle-doubled 8kHz resonant scanner with the Cousa 10×/NA0.5 objective. Calcium traces from neurons selected from the center and four corners of the field of view are plotted on the right, showing consistent signal-to-noise ratios. See also the associated **Visualization 3**.

The scan engine is compatible with nearly all commercially available objectives designed for use in multiphoton imaging applications. The beam diameter and scan angles meet or exceed the specifications of most objectives **(Fig. 5e)**. When used with a 10×/0.5 NA objective (20-mm working distance; [15]), the system achieves a 3.15 × 3.15 mm^2^ field of view **(Fig. 5f)**, and *in vivo* calcium imaging in mice demonstrated its utility **(Fig. 5g; Visualization 3)**. The resonant scanner sweeps the scan continuously across a 3.15 mm distance without stitching. This implementation showcases the generality of the angle-doubling technique. Note this version of scan-doubling actually provides for an even larger range, beyond ±5° at the objective back pupil plane. However, since we did not have access to a microscope objective that could support such wide scan angles (and even larger étendue scan optics would be required), we only used ±20° of the ±26° scan range enabled by doubling. The potential of a resonant scanned ±26° scan range, provides new design possibilities for laser scanning applications.

### 2.4 Alternative optics for angle doubling

We performed a series of modeling studies (Zemax OpticStudio; Ansys) to explore alternative optical configurations for angle doubling. The angle doubling system can be folded with a roof mirror, so that one of the relay lenses is removed and the overall size is reduced **(Fig. 6a; Visualization 4)**. In this folded configuration, the scan mirror is positioned at the front focal plane of a relay lens, and the seam of the roof mirror is positioned at the back focal plane. Based on the mirror symmetry, the collimated input beam bounced off the scan mirror, travels inside the angle doubler, and makes a loop back to the center of the scan mirror. When this collimated beam gets reflected by the same scanner the second time and leaves the system, a geometric calculation predicts that its deflection angle is doubled (*θ*_*r*_ = 2*θ*_*i*_). To confirm the doubling effect, we simulated this system with a paraxial lens, and calculated the deflection angle of the output beam (*θ*_*r*_) and the reflection angle of the input beam (*θ*_*i*_) after the scan mirror as a function the angle of the scan mirror (*α*). The result shows that the relationship of *θ*_*r*_ = 2*θ*_*i*_ holds in the simulation **(Fig. 6b; Visualization 4)**.

**Fig. 6.**
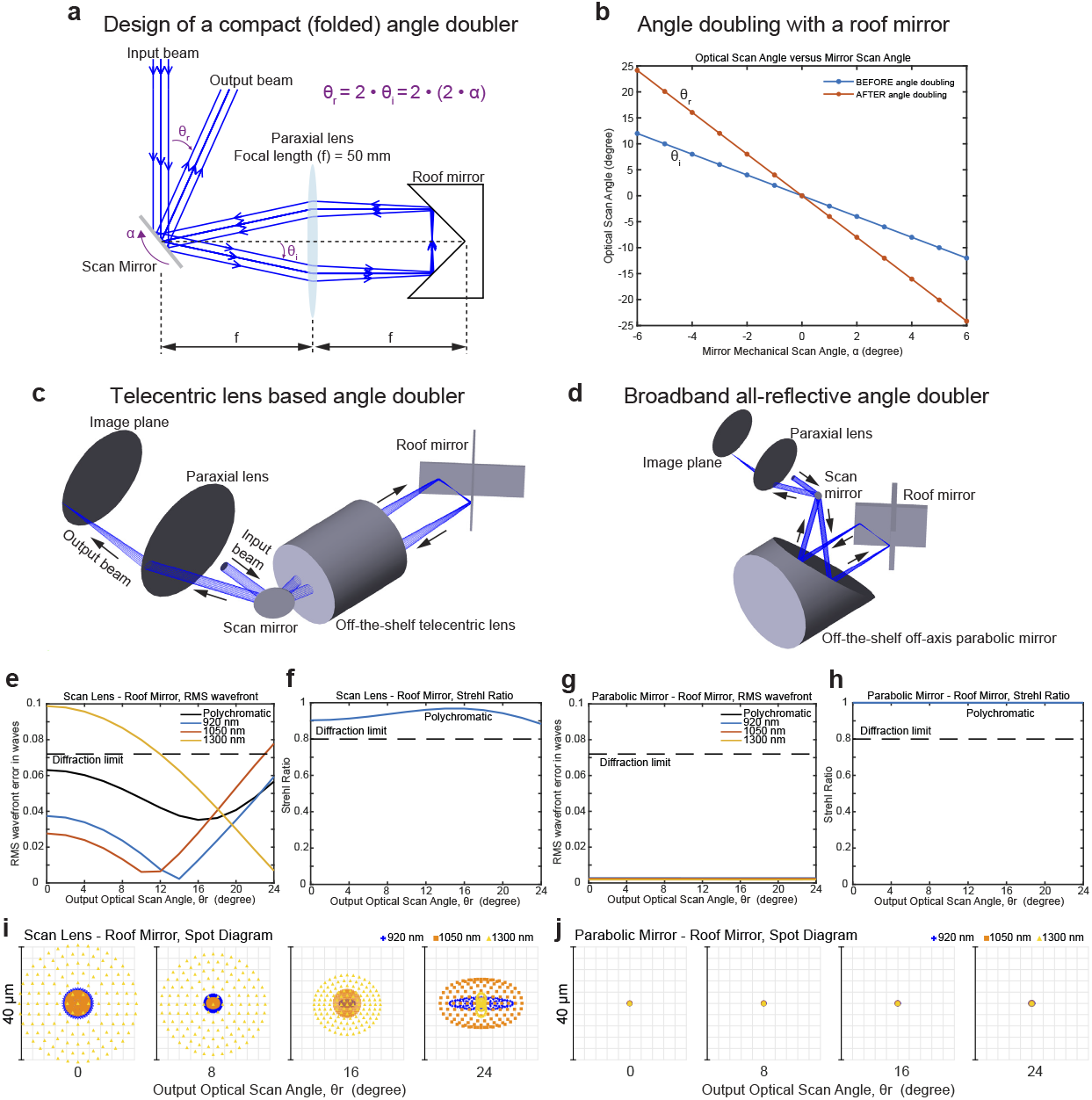
Variants of angle doublers. (a) A schematic diagram shows a folded angle doubler consisting of a paraxial lens and a roof mirror. *α* is the scan angle of the scanner relative to its neutral scan position. *θ*_*i*_ is the deflected angle of the input beam off the scan mirror relative to the optical axis before angle doubling. *θ*_*r*_ is the angle of the output beam relative to the optical axis after angle doubling. f is the focal length of the paraxial lens. See also the associated **Visualization 4**. (b) An optical simulation of the diagram (a) shows *θ*_*i*_ and *θ*_*r*_ as a function of *α. θ*_*r*_ is twice the angle of *θ*_*i*_, so angle doubling holds with this layout. (c) A schematic diagram shows a folded angle doubler constructed with an off-the-shelf telecentric lens (LSM54-1050, Thorlabs) and a roof mirror. (d) A schematic diagram shows an all-reflective angle doubler constructed with an off-the-shelf off-axis parabolic mirror (MPD249-M01, Thorlabs) and a roof mirror. See also the associated **Visualization 5**. (e) and (g) are plots of the Root-mean-square wavefront errors at 920 nm, 1050 nm, 1300 nm, and three wavelengths combined as a function of the mirror scan at the imaging plane of (c) and (d), respectively. (f) and (h) are plots of Strehl Ratio the three wavelengths combined as a function of the mirror scan at the imaging plane of (c) and (d), respectively. (i) and (j) are the spot diagrams showing the distributions of rays of 920 nm, 1050 nm, 1300 nm at the imaging plane of and (d), respectively, when the scan mirrors rotate to a different angle. Note that the final output scan angle is 24° when the scan mirror rotates to 6°.

We next replaced the paraxial lens (ideal, but unrealistic) in the model with an off-the-shelf telecentric lens (LSM54-1050; Thorlabs) and modeled the optical performance at the imaging plane **(Fig. 6c)**. We simulated the scan range of ±12° because a state-of-the-art and commonly-used resonant scanner is operating at this scan range (CRS 8 KHz, Cambridge Technology). In addition, the ±12° degree scan of the scan mirror (*θ*_*i*_) corresponds to the ±24° degree scan of the light field with angle doubling (*θ*_*r*_), which is a large scan angle compared to what current resonant scanners can offer (≤±13°, optically). The input laser is a collimated beam with a diameter of 5 mm, and contains the wavelengths of 920 nm, 1050 nm, and 1300 nm (commonly-used wavelengths for multiphoton microscopy). The root-mean-square wavefront error indicated diffraction-limited imaging for 920 nm and 1050 nm for most of the scan angle range of 0° - 12° (optical, before doubling) of the scan mirror whereas 1300 nm is not diffraction limited at small angles **(Fig. 6e)**. The result of good performance at 920 nm and 1050 nm, but not 1300 nm, is consistent with the optimization wavelength at 950 - 1150 nm for this telecentric lens. Despite some variability, the Strehl ratio of the three wavelengths combined remains diffraction-limited across 0 - 24° of the final optical scan range (*θ*_*r*_) **(Fig. 6f)**. The analysis of the ray spread (spot diagram) shows distinctive distributions of rays for different wavelengths and at different scan angles **(Fig. 6i)**. This wavelength dependence is expected, as glass is a dispersive element by nature, and performance can still be diffraction-limited over the 920 - 1050 nm wavelength range. Thus, this modeling showed that a folded (2-f) system is a feasible, with nearly half the size of a full (4-f) angle doubler and consisting of fewer components.

The wavelength dependence (chromatic aberration) can be minimized by replacing the telecentric lens with a reflective surface, such as an off-axis parabolic mirror (MPD249-M01, Thorlabs) **(Fig. 6d; Visualization 5)**. The analysis of the root-mean-square wavefront error indicates diffraction-limited performance with an extremely low wavefront error (nearly zero) at three individual wavelengths **(Fig. 6g)** and with an extremely high Strehl ratio (nearly one) for three wavelengths combined **(Fig. 6h)** over 0° - 6° scan angle (*α*). The analysis of the ray spread also shows all three wavelengths are highly concentrated, and are nearly independent of the scan angle **(Fig. 6j)**. These results demonstrate the feasibility to build a compact, all-reflective, diffraction-limited angle doubler with broadband achromaticity (wavelength-independence) and ultra-large scan angle.

The off-axis-parabolic-mirror-based angle doubler built can be further remodeled to separate the input and output beams by incorporating a resonant scanner with two reflective surfaces on both sides of the mirror substrate **(Fig. 7a; Visualization 6)**. A *ϕ*5-mm collimated beam is first reflected off one side of the resonant scanner, which scans the beam. It is then redirected to the opposite side of the scanner through a series of reflections involving six planar mirrors and two 90° off-axis parabolic mirrors, ensuring the output remains collimated. The two 90° parabolic mirrors have the same parent focal length, f (e.g., 50.8 mm), and their reflective focal lengths, f’ (e.g., 101.6 mm), is equal to 2f. This optical layout has the two parabolic mirrors to form a 4-f’ afocal optical relay with the resonant scanner. The planar mirrors are arranged such that the total beam travel distance between the resonant scanner and each parabolic mirror is 1f’ (or 2f), and the travel distance between the two parabolic mirrors is 2f’ (or 4f). When the scanning mirror rotates by *α*°, the beam undergoes a *θ*_*i*_ = 2*α*° rotation after the first reflection. Upon the second reflection off the opposite side of the scanning mirror, the output beam achieves *θ*_*r*_ = 4*α*°, realizing angle doubling. The output beam then passes through a paraxial lens with a 50.8 mm focal length, scanning a focus at an intermediate imaging plane conjugate to the focal plane under the microscope objective. Optical performance analysis at this intermediate plane demonstrates diffraction-limited performance over *θ*_*r*_ = ±24° at wavelengths of 500 nm, 920 nm, 1050 nm, 1300 nm, and 1700 nm with very low spread of ray traces **(Fig. 7b)**. The results confirm that this layout maintains the advantages of an all-reflective system, including broadband chromatic performance and an ultra-large scan angle. Additionally, the input and output beams remain spatially separated, eliminating the need for additional separation steps such as polarization optics.

**Fig. 7.**
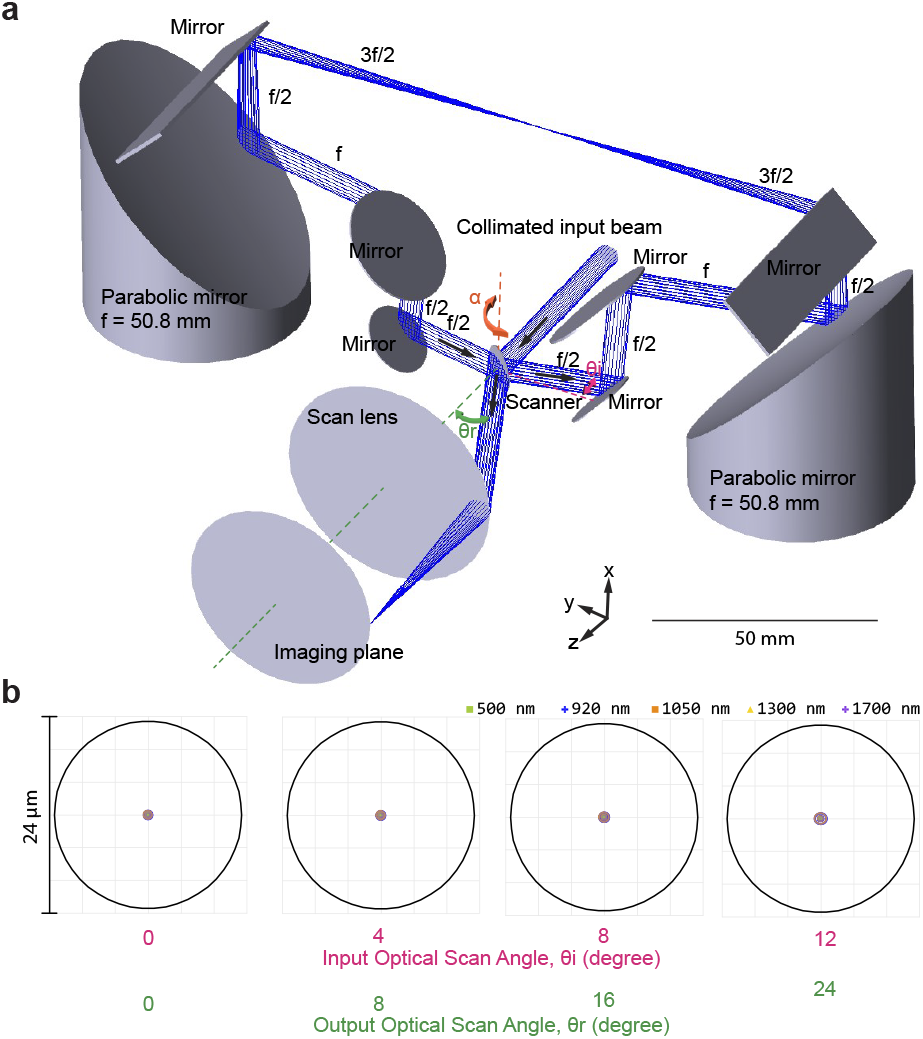
Scan angle doubler utilizing a scanning mirror with reflective surfaces on both sides. (a) Optical layout of an all-reflective angle doubler composed of a double-sided scanning mirror, 90° off-axis parabolic mirrors (MPD249-M01, Thorlabs), and planar mirrors. A 5-mm collimated input beam first reflects off one side of the resonant mirror and is then redirected to the opposite side through a system of parabolic and planar mirrors. The output scan angle, *θ*_*r*_, is two times the optical scan angle off the scanning mirror, *θ*_*i*_, achieving angle doubling. *α* represents the mirror scan angle relative to its neutral position. This diagram shows a snapshot when *α* has rotated by 6°, so *θ*_*i*_ is 12°, and *θ*_*r*_ is 24°. f represents the parent focal length of the parabolic mirror. The focal length of the scan lens is 50.8 mm. The input and output beams are spatially separated, eliminating the need for an additional step to separate them. See also the associated **Visualization 6**. (b) Spot diagrams illustrate the ray distributions for wavelengths of 500 nm, 920 nm, 1050 nm, 1300 nm, and 1700 nm at the imaging plane of (a), corresponding to different scan mirror positions. The solid circle represents the Airy disk. Across a wide range of scan angles, the rays from different wavelengths remain densely overlapped at the center of the Airy disk, demonstrating scan-angle-independent, diffraction-limited performance over an ultra-wide spectral band.

### 2.5 Phase doubling

The concept of angle doubling is extensible to phase doubling by replacing an angle modulator (a rotating mirror) with a phase modulator (such as a spatial light modulator or a deformable mirror) in a 4-f optical relay **(Fig. 8a)**. In this phase doubling system, a phase profile configured on a phase modulator is doubled in a multiplicative manner, as a flat wavefront of a collimated input beam is modulated twice for bouncing off the same modulator twice. We modeled this phase doubler (Zemax OpticStudio; Ansys) using a phase surface to mimic a 2-D spatial light modulator, and introduced a plane wave at 1300 nm in the simulation. Without phase modulation, foci are formed at the conjugated planes of A and B (gray dashed lines in **Fig. 8a**). When we applied different magnitudes of coefficients to the defocusing term of the Zernike polynomials on the phase modulator, the foci were shifted away from these two planes along the optical axis (± Δz and ±Δz’) in **Fig. 8a**. The model shows that the focal shift at plane B (±Δz’) is twice the shift at plane A (±Δz), suggesting the modulating effect on plane B is two times larger than that on plane A **(Fig. 8b). Fig. 8c** shows the defocusing wavefronts corresponding to the foci at plane A and plane B with different coefficients applied. The peak-to-valley value of the wavefront is doubled at plane B although there is only one phase modulator with a single coefficient applied. In addition, the shapes of the wavefronts at plane B with the defocus coefficient equal to 0.2 or 0.4 are the same as those at plane A with the defocus coefficient equal to 0.4 or 0.8, respectively (gray and brown highlights in **Fig. 8c**). This result demonstrates that the same effect of the phase modulation can be achieved for only half the coefficient is needed from a phase modulator paired with a phase doubler.

**Fig. 8.**
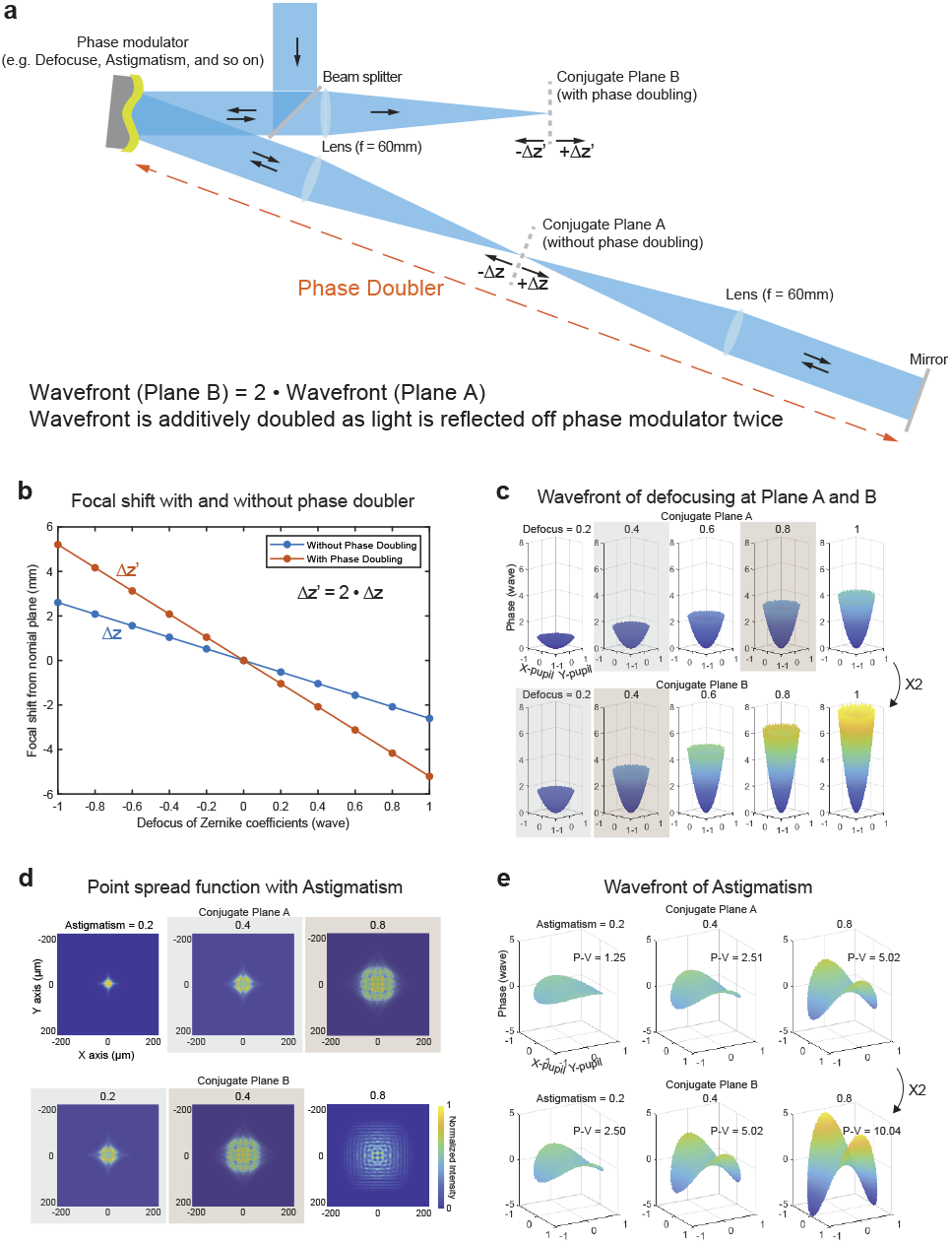
Phase doubling. (a) A schematic diagram illustrates the idea of phase doubling with a 4-f optical relay, in which a scanning mirror is replaced with a phase modulator, such as a spatial light modulator or a deformable mirror. A plane wave is launched into the phase doubler, which leads the wave to bounce off the phase modulator twice and doubles the wavefront originally applied. (b) A plot shows an optical simulation of the diagram (a) and computes the focal shift at the conjugate plane A and the conjugate plane B as a function the coefficient (0.2, 0.4, 0.6, 0.8, 1) of the defocusing Zernike polynomial applied to a 2-D phase modulator. The focal shift at the conjugate plane B is as two times large as that at the conjugate plane A. (c) The 3-D plots show the corresponding wavefront of the focus at the plane A (first row) and the plane B (second row) with different amplitudes of defocusing coefficients applied, respectively. The peak-to-valley difference of the wavefront at the plane B is as 2 times large as that at the plane A. The gray and brown colors highlight the pairs with exactly the same wavefront shapes while the amplitudes applied to the phase modulators are with a multiple of (2. d) Point spread functions at the plane A (first row) and the plane B (second row) are shown when different coefficients (0.2, 0.4, 0.8) of the astigmatism Zernike polynomial applied to a 2-D phase modulator in (a). The gray and brown colors highlight the pairs with exactly the same shapes of point-spread-functions while the coefficients applied to the phase modulators are with a multiple of 2. (e) The 3-D plots show the corresponding wavefront of the point-spread-function at the plane A (first row) and the plane B (second row) in (d), respectively. The peak-to-valley difference of the wavefront at the plane B is again as 2 times large as that at the plane A.

We next tested whether a wavefront of a non-spherical Zernike polynomial, such as astigmatism, can also be faithfully doubled in the phase doubler. When we applied different coefficients to the astigmatic Zernike polynomial on the phase modulator, we saw astigmatically aberrated point spread function at plane A and plane B. The result also shows that the shapes of the point spread functions at plane B with the coefficient equal to 0.2 or 0.4 are the same as those at plane A with the coefficient equal to 0.4 or 0.8, respectively (gray and brown highlights in **Fig. 8d**). **Fig. 8e** shows the corresponding wavefronts in **Fig. 8d**, and the peak-to-valley value of the wavefront at plane B is twice the value at plane A at each coefficient applied. These results together demonstrate that the phase doubling effect also holds for the non-spherical terms of the Zernike polynomials. Taken the results of spherical (defocusing) and non-spherical (astigmatism) wavefront modulation together, we show the feasibility to build a phase doubler using a 4-f system, and increase the dynamic range of a phase modulator by a factor of two.

### 2.6 Angle amplifying for further non-inertial increases in scan amplitude

**Figure 9a** shows a schematic representation of an inertia-free scan angle multiplier (**Visualization 7**). Like the scan doubler, it is a four focal length (4-f) optical relay with a scanning mirror and a flat mirror positioned at each side of the relay with one focal distance (f) from the nearest lens, individually. An incident beam is introduced into and reflected off a turning mirror in the space between two lenses. The reflected beam is traveling parallel to and at a distance of L from the optical axis of the 4-f relay. The principle of the angle multiplier is to retro-reflect the scanning beam off the same scanning mirror repeatedly (N times for example), and increase the maximal scan angle (Δ*θ*) upon each reflection. Based on geometric optics, the retro-reflection is realized by the arrangement of the 4-f optical relay and the flat mirror to loop the scanning beam. The flat mirror tilts from the optical axis of the 4-f relay with a fixed angle (*α* = L/(2*f*N)*(180/*π*) in degrees) that serves to separate each amplified scan from overlapping in space, and steers the plane of the final amplified scan coplanar with the optical axis before the scanning laser exits the multiplier **(Fig. 9b)**. A rectangular mirror is positioned in the middle of the optical relay to output the amplified scan with a final maximal scan angle of N*Δ*θ*. Note that this system is, like the prior arrangements in this report, a passive add-on, and can work for phase modulation as well. This system can amplify the scan angles (or phase) in an inertia-free manner, while preserving the operation of the scanner, the beam size on the scanner, and the scan frequency.

**Fig. 9.**
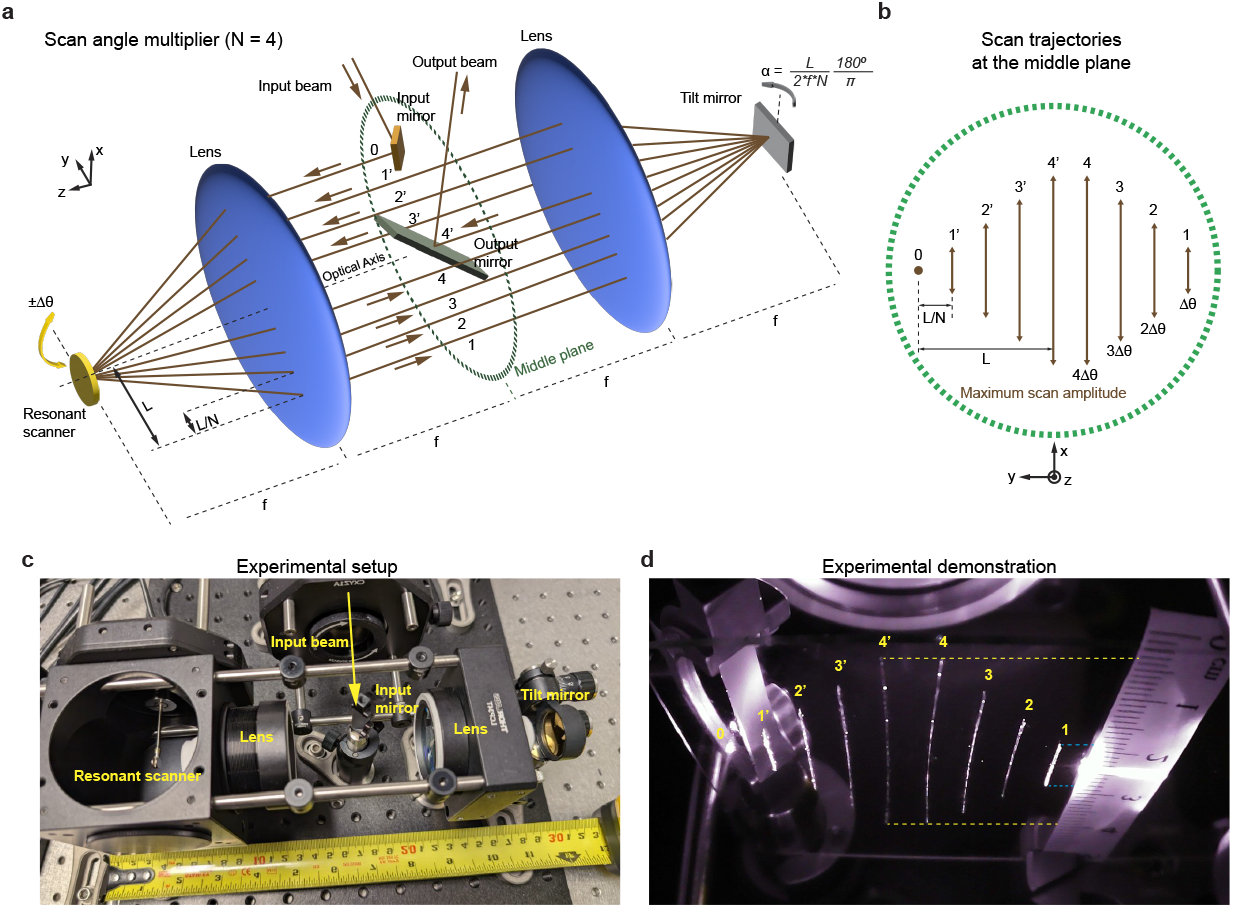
Scan angle amplifier. (a) Optical prescription of the scan angle multiplier with a multiplication factor (N) equal to 4. The system consists of a four-focal-length (4-f) optical relay incorporating a resonant scanner and a tilt mirror, each positioned on opposite sides of the relay at a focal distance (f) from the nearest lens. An incident beam enters the system and is reflected off an input mirror situated in the space between two lenses. The reflected beam propagates parallel to the optical axis of the 4-f relay at a distance of L. The resonant scanner oscillates at a scan angle of ±Δ*θ* around the y-axis. On the opposite side, the tilt mirror rotates by *α*° around the x-axis to redirect the laser beam back toward the scanner, introducing spatial separation from the previous scan. With each reflection on the resonant scanner, the scan angle increases by ±Δ*θ*. Ultimately, the amplified scan, multiplied by N, aligns with the optical axis of the relay and is extracted via a rectangular mirror. See also the associated **Visualization 7**. (b) A schematic shows an expected scan trajectories at the middle plane shown in panel (a), when the scan angle multiplier is configured for 4x amplification, with the final 4x scan traveling along the optical axis of the optical relay. The progressive amplification of the scan angle (Δ*θ*, 2Δ*θ*, 3Δ*θ*, 4Δ*θ*) is spatially separated, with scan lengths proportional to the amplification factors. (c) Photograph of the experimental setup featuring two 50-mm focal length scan lenses and a 12-kHz resonant scanner. The scan angle multiplier measures approximately one foot (∼30 cm) in length, maintaining a compact footprint. (d) Measurement of scan trajectories at the middle plane in the 4x scan angle multiplier when the 12-kHz resonant scanner operates at its maximum optical scan angle of ±5°. The observed trajectories align with the predicted paths shown in panel (b). The shortest scan (labeled 1 and 1’) measures ∼9 mm (distance between blue dashed lines), corresponding to a scan at ±5° - the scanner’s maximal capability. After 4x amplification, the longest scan (labeled 4 and 4’) extends to ∼36 mm (distance between yellow dashed lines), corresponding to a ±20° scan angle, four times larger than the original limit of the resonant scanner. The scale bar is in metric units, with ticks at one-millimeter increments. See also the associated **Visualization 8**.

To verify the concept of the scan angle multiplier, we built a proof-of-concept system **(Fig. 9c)** using a resonant scanner (Δ*θ*= ±5°, CRS 12 KHz, Novanta Optics), two telecentric lenses (Ventana SL, Pacific Optica) with the same focal length of 50 mm, and a flat mirror (PF10-03- M01, Thorlabs). The input laser was coupled into the system with a focusing lens (fi=60mm, AC254-060-B Thorlabs) and a turning mirror positioned so that L=17mm. The flat mirror was tilted at *α* =2.4°, so that the final scan was amplified by 4 times (N=4) and coplanar with the optical axis. According to the geometric optics, the length of the scanning lines at the middle plane of the system is approximately proportional to the scan angle, so we can measure the line length at that plane to infer the scan angles before and after the angle multiplication is applied. As the scan mirror scans at ±5° optically, we found that the length of the line from after amplification is (∼36 mm), and the length of line from before amplification is ∼9 mm **(Fig. 9d; Visualization 8)**. Nearly 4 times of length difference demonstrates that the scan angle is amplified by 4 times, resulting in a ±20° scan. Thus, this system amplifies the optical invariant by 4 times while the the scan frequency remains intact at 12 kHz. These results match the prediction of angle multiplication, confirming the validity of our proposed system.

## 3. Discussion

We presented an inertia-free approach to double the maximal scan angle of a galvanometer mirror scanner, while preserving its scan frequency and scan aperture. This provides performance beyond the usual trade-off between scan angle and scan aperture. In this way, the optical invariant seems not so invariant. We presented several variations, with diffraction-limited performance, including a phase doubling approach for phase modulation applications such as adaptive optics, and an angle multiplying approach, which can more than double the beam angle amplitude with no tradeoffs in beam diameter. We refer to these systems as Nisam units, for Non-Inertial Scan Angle Multipliers, specifically Nisam2x for angle doubling, Nisam4x for angle quadrupling, and so forth.

Our demonstration systems were built with resonant and linear/servo galvanometer scanners in applications for multiphoton imaging. The concept extends easily to other scan technology including micro-electromechanical systems (MEMS) scanners [19], rotating polygonal scanners, and acousto-optical deflectors (AODs). Other applications including confocal microscopy, light-sheet microscopy, light detection and ranging (LIDAR), and two-photon polymerization technology can also benefit from doubled or amplified scan angles. Many étendue-limited approaches could benefit from the technology discussed here.

Our demonstration systems used two widely used resonant scanners: a 12 kHz unit and an 8 kHz unit. Doubling the scan angle of the 12 kHz resonant scanner, from ±5° to ±10°, doubles its optical invariant, which is defined as the product of the mirror radius and the sine of the scan angle. The layout we used increased the optical invariant from 0.22 to 0.43. Next, we demonstrated angle doubling in an 8-kHz resonant scanner, resulting in an optical invariant of 1.10. This required novel scan lens and tube lens designs to accommodate the optics. The optical invariant of the 8 kHz system exceeds that of the microscope objective we used, the Cousa (10x/0.5 NA [15]), and all other infinity corrected multiphoton imaging objectives we are aware of (with the notable exception of the objective on the Diesel2p system [6]). Future objective designs could be optimized to use this scan engine to its fullest potential.

One practical implication of this technology is that it is now possible for a resonant scanner to scan the full width of a large étendue objective. High NA (> 0.5) mesoscale multiphoton systems often include resonant scan devices, but they are typically limited by their scan angle amplitude to scan only a portion of the field of view, about 0.6 mm for the 2pRAM [7] and 1.5 mm for the Diesel2p [6]. Some mesoscope systems can resonant scan the full field-of-view, but at the cost of a lower numerical aperture [5, 12]. With angle doubling or amplification, as we present here, these large field-of-view systems can provide additional imaging modes, including high speed sampling over large fields of view with no need for striping. Moreover, the angle doubler and multiplier can be compatible with other laser scanning systems, such as confocal microscopes [20], light-sheet microscopy [21, 22], harmonic and Raman-based imaging [23–27], optical coherence tomography [28–30], and advanced manufacturing with multiphoton polymerization [31].

The angle-doubling method shares similarities with the scan multiplier (SMU) [14] and the polygon scanner auto-relay [13], but it offers distinct advantages. The SMU, applied to resonant scanners, uses a micro lens array at the second lens position to divide a scan into multiple smaller ones. This multiplies the scan rate and thus enables very high update rate scanning [14, 32]. In contrast, the angle doubling and amplifying approaches we present here preserve the scan rate, and increase the scan angle, or optical invariant. The SMU arrangement divides the original optical invariant in each sub-scan, resulting in a scan angle in each sub-scan smaller than that of the original resonant scanner. Thus, these techniques have different strengths. The auto-relay technique, applied to polygon scanners [13], amplifies their optical invariant by 1.7x. This is an important advance for polygon scanners, as their optical invariants tend to be low, often less than those of resonant scan mirrors. Polygon scanners offer very fast line rates, sometimes faster than resonant scanners, and so this increase in optical invariant can be impactful across a range of applications. The angle-doubling approach we present here achieves a 2-fold improvement using resonant scanners. Thus, the SMU and the polygon scanner auto-relay provide important advances for high rate scanning. However, their optical invariants are smaller than those of standard optics and microscopes. That is a key distinction with the angle doubling and multiplying methods we present here, which provide non-inertial methods for increasing the optical invariant by 2x or larger factors.

Another novel aspect, to the best of our knowledge, is that in our angle doubler, 2D raster scanning is achieved by integrating a 1-axis linear scanner behind the second lens, scanning orthogonally (y-scan) to the resonant scanner (x-scan) within a single 4-f relay. In contrast, the polygon scanner auto-relay used an additional 4-f relay to introduce the orthogonal scanner. Thus the design we present here is compact with a small number of components, which minimizes dispersion and simplifies alignment. We also note that the single-axis linear scanner can be replaced with a dual-axis fast steering mirror. This arrangement would provide an arbitrary offset to the resonant axis (i.e., panning of the resonant scan axis across the field-of-view), thus enabling arbitrary repositioning of the field of view along both the x and y axes with no increase in components or physical footprint.

Finally, we proposed an idea of phase doubling adapted from the concept of angle doubling, and demonstrated its feasibility using optical simulations **(Fig. 8)**. Simply replacing the scanning mirror with a deformable mirror, our angle-doubling unit turns into a phase-doubling (or stroke-doubling) unit. This phase-doubling approach can benefit the application of adaptive optics. For example, a spatial light modulator for phase might have less than a full 2*π* radians of range, and angle doubling can provide an increased range. Similarly, deformable mirrors have a finite stroke or range, and the settling time scales with the stroke displacement. With phase doubling, we can increase the phase range for adaptive optics, and decouple the displacement of the stroke from the settling time with deformable mirrors. Also, the position of the deformable mirror, or any wavefront control device, is conjugate with the back aperture of the objective. This layout conforms to the standard pupil conjugation arrangement in adaptive optics, facilitating adoption of phase doubling within existing optical systems. Moreover, higher multiples are possible with phase amplification, which could be implemented by extending the scan angle amplification approach we presented.

Innovative scan technologies that are limited by étendue could benefit from scan angle doubling and amplifying, including free-space angular-chirp-enhanced delay (FACED) scanning [33–35]. Scanning with acousto-optical deflectors (AODs) is popular for random access imaging and diverse scan modes [36–43]. However, AODs typically have small scan amplitudes, and small clear apertures, since larger apertures result in slower scan speeds. In addition, novel applications of polygon mirrors can benefit from angle doubling and amplification as well [9, 13, 44–46], since these devices provide high scan rates at the cost of scan amplitude. Thus, a range of innovative laser scanning approaches can benefit from the techniques described here.

In summary, we have presented a series of non-inertial, diffraction-limited approaches for expanding scan angles and phase modulation, from doubling to amplification to an arbitrary range. The net result is an increased optical invariant, which can benefit a range of applications from laser-based imaging, as we demonstrate here, to adaptive optics, LIDAR, lightsheet imaging, lithography, manufacturing, other laser scanning and wavefront modulation approaches.

## Supporting information

Visualization 1

Visualization 2

Visualization 3

Visualization 4

Visualization 5

Visualization 6

Visualization 7

Visualization 8

## Acknowledgments

The authors thank I. Smith, F. Tomaska, and members of the lab for valuable discussions and feedback.

## Data availability

Data underlying the results presented in this paper are not publicly available at this time but may be obtained from the authors upon reasonable request

### Visualizations Visualization 1

In vivo calcium imaging (GCaMP6s) in a mouse using the angle-doubled 12kHz resonant scanner and the Nikon 16x water immersion objective (0.8 NA). FOV: 1.05 × 1.05 mm^2^. Frame rate: 45 frames/s. Pixel size: 512 × 512 pixels. Power at the imaging plane: 30 mW. Play Speed: 5x.

**Visualization 2**

In vivo calcium imaging (GCaMP6s) in a mouse using the angle-doubled 12kHz resonant scanner and the Cousa 10x air objective (0.5 NA, 20mm working distance). FOV: 1.7 × 1.7 mm^2^. Frame rate: 15.6 frames/s. Pixel size: 1536 × 1536 pixels. Power at the imaging plane: 50 mW. Play Speed: 10x.

**Visualization 3**

In vivo calcium imaging (GCaMP6s) in a mouse using the angle-doubled 8kHz resonant scanner and the Cousa 10x air objective (0.5 NA, 20 mm working distance). FOV: 3.15 × 3.15 mm^2^. Frame rate: 7.68 frames/s. Pixel size: 2048 × 2048 pixels. Power at the imaging plane: 70 mW. Play Speed: 10x.

**Visualization 4**

An animation shows an angle doubler using a paraxial lens and a roof mirror. *α* is the scan angle of the scanner relative to its neutral scan position. *θ*_*i*_ is the deflected angle of the input beam off the scan mirror relative to the optical axis before angle doubling. *θ*_*r*_ is the angle of the output beam relative to the optical axis after angle doubling.

**Visualization 5**

An animation shows an all-reflective angle doubler using an off-axis parabolic mirror and a roof mirror.

**Visualization 6**

An animation shows an all-reflective angle doubler using a double-sided scanning mirror, off-axis parabolic mirrors, and planar mirrors.

**Visualization 7**

An animation shows the optical description of a scan angle multiplier with a multiplication factor equal to 4.

**Visualization 8**

The video demonstrates the amplified scan trajectories at the middle plane in the 4× scan angle multiplier, as a 12-kHz resonant scanner expands its optical scan angle from 0 to ±5°.

